# Reply to Lazaridis and Reich: robust model-based inference of male-biased admixture during Bronze Age migration from the Pontic-Caspian steppe

**DOI:** 10.1101/122218

**Authors:** Amy Goldberg, Torsten Günther, Noah A. Rosenberg, Mattias Jakobsson

## Abstract

Comparing the sex-specifically inherited X chromosome to the autosomes in ancient genetic samples, we (*1*) studied sex-specific admixture for two prehistoric migrations. For each migration, we used several admixture estimation procedures—including *Admixture* model-based clustering (*2*)—comparing X-chromosomal and autosomal ancestry in contemporaneous Central Europeans, and interpreting greater admixture from the migrating population on the autosomes as male-biased migration. For migration into late Neolithic/<u>B</u>ronze <u>A</u>ge Central Europeans (“BA”) from the Pontic-Caspian steppe (“SP”), we inferred male-biased admixture at 5-14 males per migrating female.

Lazaridis & Reich (*3*) contest this male-biased migration claim. For simulated individuals, they claim that *Admixture* provides biased X-chromosomal ancestry estimates. They argue that if the bias is taken into account, then X-chromosomal steppe ancestry is similar to our autosomal ancestry estimate, and that hence, steppe male and female contributions are similar.

Many factors affect ancestry inferences from *Admixture* and related programs (*2*, *4*-*8*). To understand *Admixture* inferences for X-chromosomal ancient DNA, we performed simulations examining the effects of multiple variables. First, we used “reference” individuals in (*1*) to simulate analogs of the BA population.

Figure 1 plots estimated X-chromosomal ancestry for simulated BA individuals (Fig. 1A,B), showing that for high true ancestry levels, *Admixture* overestimates steppe ancestry, whereas for low levels, it underestimates it. For the intermediate ancestry in (*1*) (0.366), however, *Admixture* is accurate, and our estimate is robust to bias.

As our interest in (*1*) was the X/autosomal comparison, we next simulated autosomes, finding bias similar to the X chromosome (Fig. 1C,D). Bias-corrected X/autosomal ancestry estimates translate in a constant-admixture model (*1*) to 4-7 migrating steppe males per female. Thus, accounting for *Admixture* bias, substantial male excess during the steppe migration remains supported.

**Figure 1.**
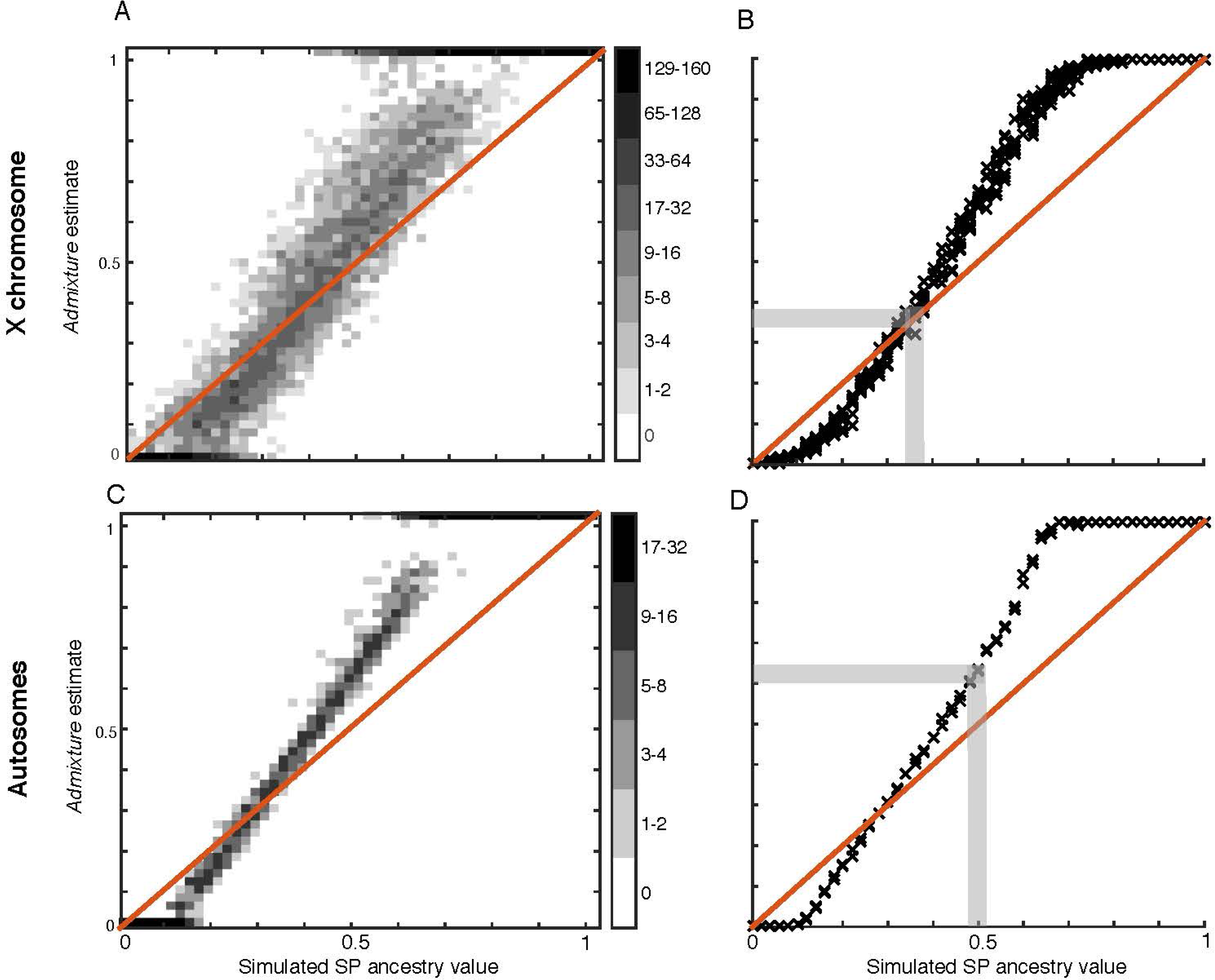
*Admixture* inference in simulated ancient genomes. We simulated admixed BA individuals for steppe-related ancestry values in 0.02 increments. Using LD-pruned SNP sets from (*1*), independently at each SNP, we drew a reference population and chose the allele randomly from among individuals in that population. We used a 10-seed average with supervised *Admixture* (*1*), considering replicate 16-individual populations. Shaded bars (panels B, D) show the range of simulated ancestries that corresponds to estimated ancestries in (*1*): [0.34, 0.38] for the X, corresponding to the same range in simulated values, and [0.60,0.64] for autosomes, corresponding to ∼0.500. The updated autsomal ancestry value generates X/autosomal ancestry ratio 0.366/0.500=0.732, compared with 0.592 in (*1*); this ratio generates an inference of 4-7 migrating males per female by the mechanistic model of (*1*).

We next tested if specific data features—haploid ancient genotypes, high missing-data rates, and small reference samples—might underlie previously unseen *Admixture* biases. We performed analogous simulations using modern HapMap samples without these features. This analysis traces the bias to the small reference samples available in haploid ancient data (*1*) (Fig. 2).

**Figure 2.**
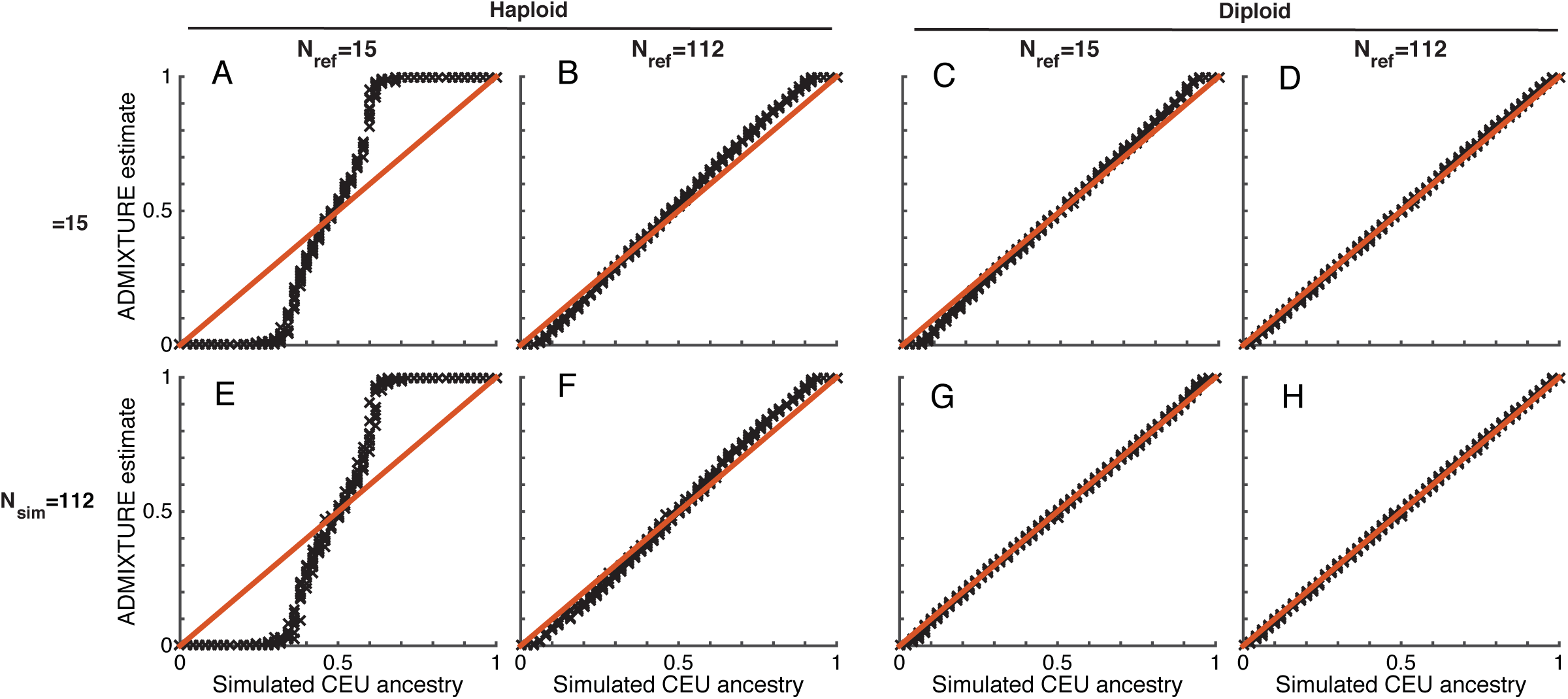
*Admixture* sample-size effects. Following the procedure in Fig. 1, we estimated ancestry in simulated admixed haploid and diploid genomes. Considering 4,605 randomly sampled autosomal SNPs, we varied the HapMap YRI and CEU sample sizes used per source population when simulating genomes (*N*_sim_) and as reference samples in *Admixture* (*N*_ref_). With Nref comparable to (*1*), *Admixture* bias in haploid simulations matches that observed for ancient data (panel A). Therefore, the bias is likely not due to missingness. As *N*_ref_ increases (B,F), bias decreases. Using differing *N*_sim_ and *N*_ref_, we attribute the bias to small reference samples during inference rather than simulation (B vs. E). Diploid simulations have minimal bias (C,D,G,H), suggesting that haploidy of ancient data combines with small reference sizes to generate the observed bias in *Admixture*.

The greater bias in *Admixture* in (*3*) than here thus likely arises from two sources. First, ancestry values underlying the simulation in (*3*) trend toward parameter values that generate higher bias than with our even spacing. Second, *Admixture* inference in (*3*) discards one individual per source population, potentially enlarging bias from small reference samples.

We note that (*3*) also consider a second program, *qpAdm* (*9*); their wide confidence intervals for this summary-statistic method relying on *f*4 calculations permit multiple interpretations (male bias, female bias, no bias). Direct *f*4 calculation (*10*), however, trends toward male-biased migration: BA share more alleles with SP than with early Neolithic Central Europeans (CE) on autosomes (*f*4(Chimp,BA;CE,SP)=0.0014; *Z*=6.78, *P*<0.0001), but have more CE X-chromosomal sharing (*f*4=−0.0068; *Z*=-0.561).

We conclude that our inference of male-biased Pontic-Caspian steppe migration, seen using *Admixture, Structure*, mechanistic simulations, and X/autosomal *F_st_*, is robust. Our analysis further illuminates the impact of small haploid reference samples on *Admixture*; we look forward to refining sex-specific migration estimates as larger, higher-coverage ancient samples become available.

## Acknowledgements

We thank Jonathan Kang and Joshua Schraiber for helpful discussions.

